# Targeted Depletion of rRNAs and Tobamoviruses Enhances Viral Discovery in Wastewater Metatranscriptomes

**DOI:** 10.64898/2026.10.08.757481

**Authors:** María A. Bautista, Tulika Bhardwaj, Dylan J. Barbera, Samuel J. Rulli, Jackie Zorz, Janine McCalder, Chloe Papparis, Xiaoli Pang, Bonita E. Lee, Judy Qiu, Rhonda G. Clark, Nicole Acosta, Christine O’ Grady, Michael D. Parkins, Casey R.J. Hubert

**Author notes:** **CRediT author statement** **María A. Bautista:** Conceptualization, Methodology, Investigation, Visualization, Writing – Original Draft; **Tulika Bhardwaj:** Software, Formal analysis, Data Curation, Writing – Review & Editing; **Dylan J. Barbera:** Methodology, Writing – Review & Editing; **Samuel J. Rulli:** Methodology, Writing – Review & Editing; **Jackie Zorz:** Visualization, Writing – Review & Editing; **Janine McCalder:** Investigation; **Chloe Papparis:** Investigation; **Xiaoli Pang:** Funding acquisition, Writing – Review & Editing; **Bonita Lee:** Funding acquisition, Writing – Review & Editing; **Judy Qiu:** Funding acquisition, Writing – Review & Editing; **Rhonda G. Clark:** Project administration; **Nicole Acosta:** Project administration; **Christine O’ Grady:** Funding acquisition; **Michael D. Parkins:** Funding acquisition, Writing – Review & Editing; **Casey R.J. Hubert:** Conceptualization, Supervision, Writing – Review & Editing, Funding acquisition.

## Abstract

Wastewater contains a complex mixture of nucleic acids derived from human cells, microorganisms, diets, and agricultural or industrial inputs. Metatranscriptomics has the potential to broaden the understanding of wastewater viromes, but its utility is constrained by high-abundance RNA molecules that prevent the detection of lower-abundance signals, including human pathogens. Two major sources of this interfering background signal are ribosomal RNAs (rRNAs) from various organisms, and plant-infecting tobamoviruses from human diets, which are often the most abundant RNA viruses in wastewater. These rRNA molecules and RNA genomes can account for upwards of 95% of total RNA sequences, thereby obscuring the detection of lower-abundance viruses of public health importance. This study evaluated whether combined depletion of ribosomal RNAs and genomic RNA from abundant plant tobamoviruses can improve the recovery of underrepresented viral sequences from wastewater. RNA extracts from a large municipal wastewater treatment plant in Calgary, Canada, were processed with or without probe-based depletion of rRNAs and eight abundant tobamoviruses. Metatranscriptomes revealed that depletion reduced rRNA reads by 90%. Reads assigned to tobamoviruses, which accounted for up to 25% of viral reads in untreated samples, were reduced to <2% of viral reads after depletion. This decrease in undesired RNA signal led to an order of magnitude increase in viral read signals and improved assembled viral genome quality. Furthermore, the number of reads assigned to human and animal pathogens increased 42-fold after depletion, enabling the detection of important pathogens like influenza, RSV, and hepatitis A, which were undecteable in untreated samples. Together, these findings demonstrate that targeted depletion of abundant background RNAs improves the detection of low abundance viruses, and enhances the utility of metatranscriptomic approaches for wastewater viral surveillance.

## Introduction

Wastewater based surveillance (WBS) in increasingly being used as a tool to provide a population-level view of the circulating viruses in a community. As a complement to traditional clinical surveillance, wastewater can be used as an unbiased record of human pathogens, independent of the symptoms and health-seeking behaviour of the individuals in a community. Widely adopted during the COVID-19 pandemic, WBS has been applied to identify and track SARS-CoV-2 and its variants^1–5^ and to support the detection of other viruses of public-health interest ^6–10^. The global use of WBS illustrates the utility of this strategy for monitoring and detecting endemic and emerging viral threats^11–13^.

WBS can be PCR based, where specific viral pathogens are detected through targeted molecular methods ^14,15^, or sequencing based, where shotgun sequencing is used to characterize the genetic material of the entire sample ^8,16–18^. While PCR based methods can be highly sensitive, cheaper, faster to perform, and easily integrated with clinical testing, sequencing based WBS offers the added benefit of capturing a wider range of pathogens that may be of public health concern but are not consistently screened for (e.g., avian flu and measles) ^6,7,11^. However, human viral pathogens only account for an exceedingly small fraction of wastewater genetic material (generally <0.1%) ^19^, which complicates sequencing based approaches. Consequently, methods including ultra-deep shotgun sequencing, on the order of billions of reads per sample^17^, or targeted hybrid capture panels that incorporate millions of sequence enrichment probes, ^8,20^ have been proposed. Both of these strategies have inherent obstacles to widespread use. Ultra-deep shotgun sequencing is expensive and has a massive computational burden that is generally not feasible for most research or clinical programs, and using thousands of viral enrichment probes can complicate wet lab procedures while significantly adding to laboratory costs^21^.

Sequencing based strategies for WBS are further challenged by abundant background RNA. For instance, ribosomal RNAs (rRNAs) comprised a large proportion (as much as 95%) of total reads in an ultra-deeply sequenced metatranscriptome ^22^. And even among the detected viral reads, a few specific plant pathogens, like the Tomato Brown Rugose Fruit Virus and other tobamoviruses, dominated the signal^23^. Tobamoviruses enter wastewater through dietary consumption of infected plant material, food-processing waste, or agricultural inputs to the sewershed, and their high relative abundance can dominate wastewater sequencing data, masking signals from less abundant pathogens of greater relevance to public health surveillance.

Previously, efforts to improve the efficiency of sequencing based WBS have focused on viral concentration in samples, viral genomic extraction, and targeted enrichment strategies ^24^. A complementary strategy is the reduction of these abundant but non-priority RNA sequences using depletion probes before library preparation to increase the proportion of genomic material from pathogens of interest in the sequencing pool^19^. Such an approach could increase the proportion of sequencing reads for broad virome characterization while retaining the pathogen-agnostic advantages of untargeted sequencing.

Here we evaluate a combined depletion strategy targeting two dominant sources of background RNA in wastewater metatranscriptomes: ribosomal RNAs and eight prevalent tobamoviruses. We compared RNA libraries prepared with and without the depletion of these RNAs to determine whether targeted subtractive removal improves the recovery of non-tobamovirus viral reads and assembled viral contigs. Using three composite influent wastewater samples collected from a large municipal treatment facility in Calgary, Canada, we demonstrate that this approach increases the relative abundance of otherwise underrepresented wastewater RNA viruses in metatranscriptomic libraries.

## Materials and Methods

### Sample collection and pre-processing

Wastewater was collected as flow-weighted 24-hour composite samples from Calgary’s principal wastewater treatment plant (WWTP) serving a catchment of ∼1.04 million individuals. Sampling used ISCO 6212 or ISCO 5800 (Teledyne ISCO, USA) portable autosamplers as previously described ^25^. Calgary’s wastewater system collects discharges from residential, commercial, and industrial buildings, with negligible agricultural input. Stormwater is managed separately, reducing the influence of rainfall on the wastewater sampled at the treatment plant. Once collected, samples were stored at 4°C for 2–4 hours before being transported to the University of Calgary’s Advancing Canadian Water Assets (ACWA) laboratory for pre-processing. Sampling took place every two weeks on May 14, 2024 (WW-1), May 28, 2024 (WW-2) and June 11, 2024 (WW-3). Each sample consisted of 360 mL of raw wastewater split evenly into eight individual 50 mL conical tubes, each containing 45 mL of raw wastewater. Tubes were centrifuged at 4,000 × g for 30 min at 4 °C to remove solids. Supernatant was filtered through a 5 μm PVDF (MilliporeSigma, USA) membrane to remove any remaining large particles that did not pellet during centrifugation. This resulted in 320 mL of total filtered supernatant that was re-pooled then concentrated using ultrafiltration with a 30 kDa Centricon® Plus centrifugal filter unit (MilliporeSigma, USA), reducing the volume down to approximately 40 mL. Concentrated supernatant was immediately placed on ice and transported to the Geomicrobiology laboratory on the University of Calgary campus, where RNA extraction was performed immediately upon arrival.

### Nucleic acid extraction

Each 40 mL concentrated wastewater filtrate was split into two 20 mL aliquots and processed using the MagMAX™ Wastewater Ultra Nucleic Acid Isolation Kit with Virus Enrichment (Applied Biosystems™, USA). The manufacturer’s instructions were modified slightly by doubling the reaction volume and each 20 mL aliquot had 200 μL of Dynabeads™ Wastewater Virus Enrichment beads (Applied Biosystems™, USA) added. Nucleic acids from each 20 mL aliquot were eluted in 60 μL of Nuclease-Free Water (QIAGEN, Germany), and the two eluates from each sample were pooled together prior to DNase treatment using the rigorous protocol recommended for the Turbo DNA-free™ Kit (Invitrogen, USA), with a total of 6 U of TURBO DNase per reaction. For every 45 μL of sample, 0.1 volume of 10× TURBO DNase™ Buffer and 3 U of TURBO DNase were added to the extracted nucleic acids and incubated for 30 min at 37°C. An additional 3 U of TURBO DNase were then added, followed by a 30 min incubation at 37 °C. Tubes were gently inverted by hand every 15 min during both incubations. Finally, 0.2 volumes of DNase Inactivation Reagent was added. After centrifugation, all fractions of the same sample were pooled into a new 1.5 mL low-retention tube, thoroughly mixed then split into aliquots, and immediately stored at −80 °C until further processing.

### Custom tobamovirus probe design

QIAseq FastSelect™ Custom RNA removal probes (available from Qiagen as FastSelect Panel ID: CTFS-05111Z-369) (QIAGEN, Germany) were designed for this study, to target eight of the most abundant tobamoviruses found in wastewater metatrascriptomes ^26,27^. QIAseq FastSelect probes are designed to prevent cDNA synthesis of the target sequences during reverse transcription. The eight viruses targeted for probe-based removal in this way were: Tomato Brown Rugose Fruit Virus (NC_028478.1), Pepper Mild Mottle Virus (NC_003630.1), Tobacco Mild Green Mosaic Virus (NC_001556.1), Cucumber Green Mottle Mosaic Virus (NC_001801.1), Tomato Mosaic Virus (NC_002692.1), Bell Pepper Mottle Tobamovirus (NC_009642.1), Tomato Mottle Mosaic Virus (NC_022230.1), Tropical Soda Apple Mosaic Virus (NC_030229.1).

### Tobamovirus and ribosomal RNA depletion, library preparation, and sequencing

To compare the effect of rRNA and tobamovirus genomic RNA depletion on viral read output in wastewater metatranscriptomes, one aliquot from each of sample was treated with FastSelect depletion probes (hereafter referred to as treated), whereas a second parallel aliquot entered the workflow at the cDNA synthesis stage without depletion (untreated). This generated a total of six metatranscriptome libraries.

Simultaneous removal of bacterial rRNA and tobamovirus RNA was achieved by combining QIAseq® FastSelect™–5S/16S/23S kit (QIAGEN, Germany) with the tobamovirus RNA custom panel following manufacturer’s instructions for QIAseq® FastSelect™ probes. To the initial 11.5 μL RNA sample, 1 μL of FastSelect 5S/16S/23S, 1 μL of the custom tobamovirus depletion probes, and 1.5 μL of FastSelect FH Buffer were added. The reaction mixtures were then incubated according to the profile recommended for degraded RNA (RIN ≤ 2): 2 min at 75°C, 2 min at 70°C, 2 min at 65°C, 2 min at 60°C, 2 min at 55°C, 2 min at 37°C, 2 min at 25°C, with a 4°C hold at the end. After bead cleanup, RNA was eluted in 12 μL of nuclease-free water.

For cDNA synthesis with the QIAseq xHYB Viral RT Kit (QIAGEN, Germany) the maximum allowed input (5 μL) was used. Template for cDNA synthesis was either RNA treated with depletion probes or untreated RNA. Elution of cDNA in 18.25 μL nuclease-free water following the recommended bead cleanup steps, and 15.75 μL was transferred to a fresh tube to avoid bead carryover. Library preparation then proceeded using the QIAseq FX Kit (QIAGEN, Germany). For the FX reaction, all 15.75 μL of cDNA was used, following kit instructions for samples with <10 ng input DNA, including the use of enhancer. QIAseq UDI Y-Adapters were diluted 1:10, and libraries were amplified with 12 PCR cycles. Final libraries were eluted in 30 μL of nuclease-free water then quantified on a Qubit™ 4 Fluorometer (Invitrogen, USA) using the Qubit™ 1X dsDNA High Sensitivity Kit (Invitrogen, USA). Library quality control was performed on an Agilent® 2100 Bioanalyzer with the High Sensitivity DNA Analysis Kit (Agilent Technologies, USA). Shotgun sequencing of the six libraries was carried out by Novogene Corporation Inc. (USA) on a full lane of an Illumina NovaSeq X 25B flow cell using a paired-end 2×150 bp strategy. The six metatranscriptome libraries contained on-average 535,442,635 reads.

### Metatranscriptome pre-processing

Raw metatranscriptomic datasets (FASTQ format) were checked for data quality and processed which included initial quality assessment and fastp (v0.24.0) for adapter trimming and quality filtering with default parameters and removal of reads shorter than 50 bp, before downstream analyses using FastQC (v0.11.9). For accurate and rapid ribosomal RNA sequence detection based on deep learning, RiboDetector (v0.2.7) was used in paired-end mode ^28^. It employed pre-trained deep learning models derived from curated rRNA databases (i.e., SILVA and RDP Databases). Reads were classified as rRNA, non-rRNA, or unclassified based on alignment to curated rRNA reference databases. Only non-rRNA read pairs were retained for downstream viral analyses. A combined identity threshold of >95% and coverage threshold (i.e., the minimum proportion of the rRNA sequence that a read must cover to be classified as rRNA) of >75% were applied. RiboDetector was executed with an expected read length of 100 bp (-l) to ensure optimal model performance, a chunk size of 256 (-c) to balance computational efficiency and memory usage during batch processing, and 20 CPU threads (-t) to enable parallel execution of the computation at High-Performance Cluster (HPC).

### Read-based viral profiling

Kraken2 ^29^ was used for taxonomic classification of non-rRNA read pairs against a curated viral reference database constructed from NCBI RefSeq viral genomes (downloaded in September 2024), and reads were assigned based on exact k-mer matches using a lowest common ancestor (LCA) algorithm with default parameters. Viral taxonomic profiles were summarized using Kraken2 report files, followed by calculating Bayesian re-estimation of the abundance of Kraken reads using Bracken (v2.9) ^30^. Classification was performed using a k-mer length of 35 (-- kmer-len), a minimizer length of 31 (--minimizer-len), and 7 minimizer spaces (--minimizer-spaces). A confidence threshold of 0 requiring at least 2 distinct minimizer hit groups (--minimum-hit-groups) and a minimum base Phred quality of 0 (--minimum-base-quality) was used.

To specifically identify viral pathogens of humans and animals, EsViritu (v1.3.3) ^8,31^ was employed on the paired-end non-rRNA reads. This reference-based viral detection tool maps sequencing reads against a curated database (EsViritu database v3.2.4) of human and animal viral genomes, clustered at approximately 95% nucleotide identity. After alignment, virus detection was performed based on sequence identity and genome coverage at default settings. This enables highly sensitive identification of human-, plant- and animal-associated viral lineages in wastewater, and complements Kraken2 taxonomic profiling by providing higher resolution for human and animal-pathogen lineages. As Tobamoviruses are included in the EsViritu database and were targeted by the depletion treatment, reads assigned to the Tobamovirus genus were excluded from downstream analyses to prevent the depletion procedure from confounding comparisons of viral abundance.

### Reference-based mapping for targeted tobamovirus quantification

To determine specific tobamoviruses, non-rRNA reads were mapped to eight reference tobamovirus genomes using BBMap (v38.22) ^32^, with default alignment parameters, generating SAM-formatted alignments. Resulting SAM files were converted to BAM format, sorted and indexed using SAMtools (v1.20) to facilitate efficient downstream processing (Li et al., 2009). Alignment statistics were then summarized using SAMtools flagstat, including the proportion of mapped reads.

### Contig-based viral discovery and genome recovery

Non-rRNA reads were assembled *de novo* with MEGAHIT (v1.2.9), which employed a succinct de Bruijn graph framework ^33^, using default parameters. Assembled contigs were checked for viral sequences using geNomad (v1.8.1) ^34^, a high-throughput platform that utilizes machine learning-based models trained on sequence comparison, gene content, and marker features to distinguish viral and plasmid contigs from other genomic sequences. Contigs were identified as viral or plasmid-derived based on a minimum score threshold of 0.7 for both types. Only contigs with a minimum length of 1,000 bp and at least one predicted gene were included. Viral contigs identified by geNomad were assessed for quality using CheckV (v1.0.3) ^35^, based on the estimated genome completeness by comparing them to the curated Global Viral Database (GVDB). Based on CheckV quality tiering, viral contigs were categorized as high-quality (>90% completeness), medium-quality (50-90% completeness), and low-quality (<50% completeness).

All analyses were implemented as a fully reproducible and modular workflow using Nextflow ^36^. The workflow was executed on the Advanced Research Computing (ARC) HPC cluster at the University of Calgary, ensuring robustness and reproducibility in different computing clusters. Containerization using Singularity was employed to maintain consistent software environments across runs. The workflow is publicly available at https://github.com/tulikasriv26/MetavIRAL-.

## Results

### Ribosomal RNA and Tobamovirus RNA depletion enables enhanced viral read recovery

The fraction of reads annotated as rRNA decreased from 92–93% to <5% in the rRNA and Tobamovirus depleted samples (treated) compared to the controls (untreated) (Figure 1A, Supplementary Table 1). In untreated samples, only 3–4% of reads were classified as non-rRNA, whereas treated samples showed a corresponding increase in non-rRNA reads to 94–96%. Overall, rRNA depletion resulted in an approximate 20–25-fold increase of non-rRNA sequences, substantially expanding the proportion of reads suitable for downstream metatranscriptomic analysis of RNA viruses. A small fraction of reads in the treated libraries remained unclassified (1-4%), likely reflecting sequences with insufficient discriminatory signal for confident classification.

**Figure 1.**
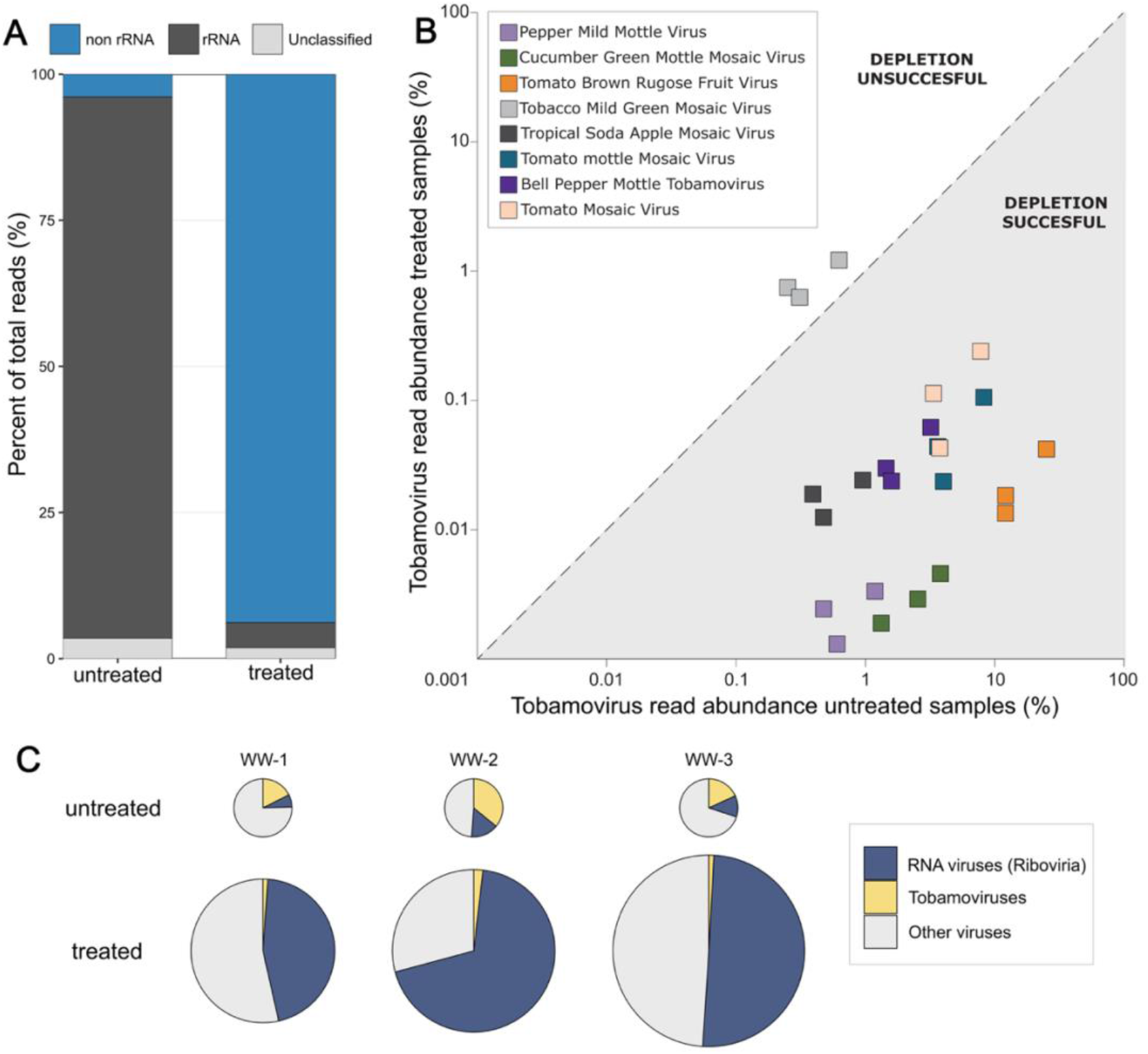
Effects of rRNA and tobamovirus depletion on read composition and viral community profiles in wastewater metatranscriptomes. (A) Average fraction of raw reads assigned to rRNA, non rRNA, and unclassified categories using RiboDetector, with (treated) and without (untreated) rRNA and Tobamovirus depletion probes. (B) Tobamovirus abundance in untreated (x-axis) versus treated (y-axis) libraries using each of the eight depletion probes. The dotted line denotes no change; the grey shaded region indicates successful depletion (Treated < Untreated). Axes have been log-transformed. (C) Distribution of viral reads in treated and untreated samples. The size of the pie chart in the treated sample is proportional to the increase in the overall number of Kraken2-classified viral reads compared to its corresponding untreated sample. Treated samples had 6.2x, 7.9x, and 10.9x more Kraken2-classified viral reads than their untreated counterpart for WW-1, WW-2, and WW-3 respectively.

Genome-specific mapping of raw reads to the eight tobamovirus genomes used to design the probe panel showed marked reductions in these tobamovirus signals after depletion (Figure 1B, Supplementary Table 2). The amount of viral classified reads mapping to Tomato Brown Rugose Fruit Virus, the most abundant tobamovirus in untreated samples, decreased from 12% to 0.02% in WW-1, from 25% to 0.04% in WW-2, and from 12% to 0.01% in WW-3. As shown in Figure 1B, a decrease in reads mapping to each tobamovirus was observed in all depleted samples except for Tobacco Mild Green Mosaic Virus, for which the number of mapped reads slightly increased.

The number of Kraken-2 classified viral reads increased by 6- to 11-fold after depletion treatment of the three wastewater samples, reaching up to 4.8 million reads (Figure 1C, Supplementary Table 1). In untreated samples, the fraction of reads classified as viral by Kraken2 was very low, ranging from 0.09-0.15%, while in treated samples viral classified reads increased to 0.9-1.2% (Supplementary Table 1). Additionally, RNA viruses (Riboviria), and in particular the non-Tobamovirus fraction, increased from, 11.4% to 54.9% of viral reads on average (Figure 1C). This highlights that both the absolute quantity and total proportion of RNA viruses of interest increased after the depletion treatment.

### Assembly-based viral genome recovery

While read-based analysis of wastewater metatranscriptomic sequences is generally sufficient for the surveillance of viral pathogens, assembling viral reads into longer contigs can provide additional genomic context, enabling the detection of divergent or previously uncharacterized viruses and improving discrimination among closely related viral variants and phylogenetic lineages. Here, the number of assembled viral contigs increased in all samples treated with depletion probes. Depletion libraries resulted in more viral contigs with at least one hallmark gene. Numbers of viral contigs of 5,450, 10,356, and 5,708 in the three untreated samples increased to 46,328, 57,068, and 53,083 following depletion (Figure 2, Supplementary Table 3) for WW-1, WW-2, and WW-3, respectively. This corresponds to increases in treated samples of 5.5 to 9.2 fold over untreated samples (Figure 2). Most of these viral contigs belonged to the phylum Lenarviricota (mainly bacteriophages), with the phyla Kitrinoviricota and Pisuviricota (containing human pathogens) also represented.

**Figure 2.**
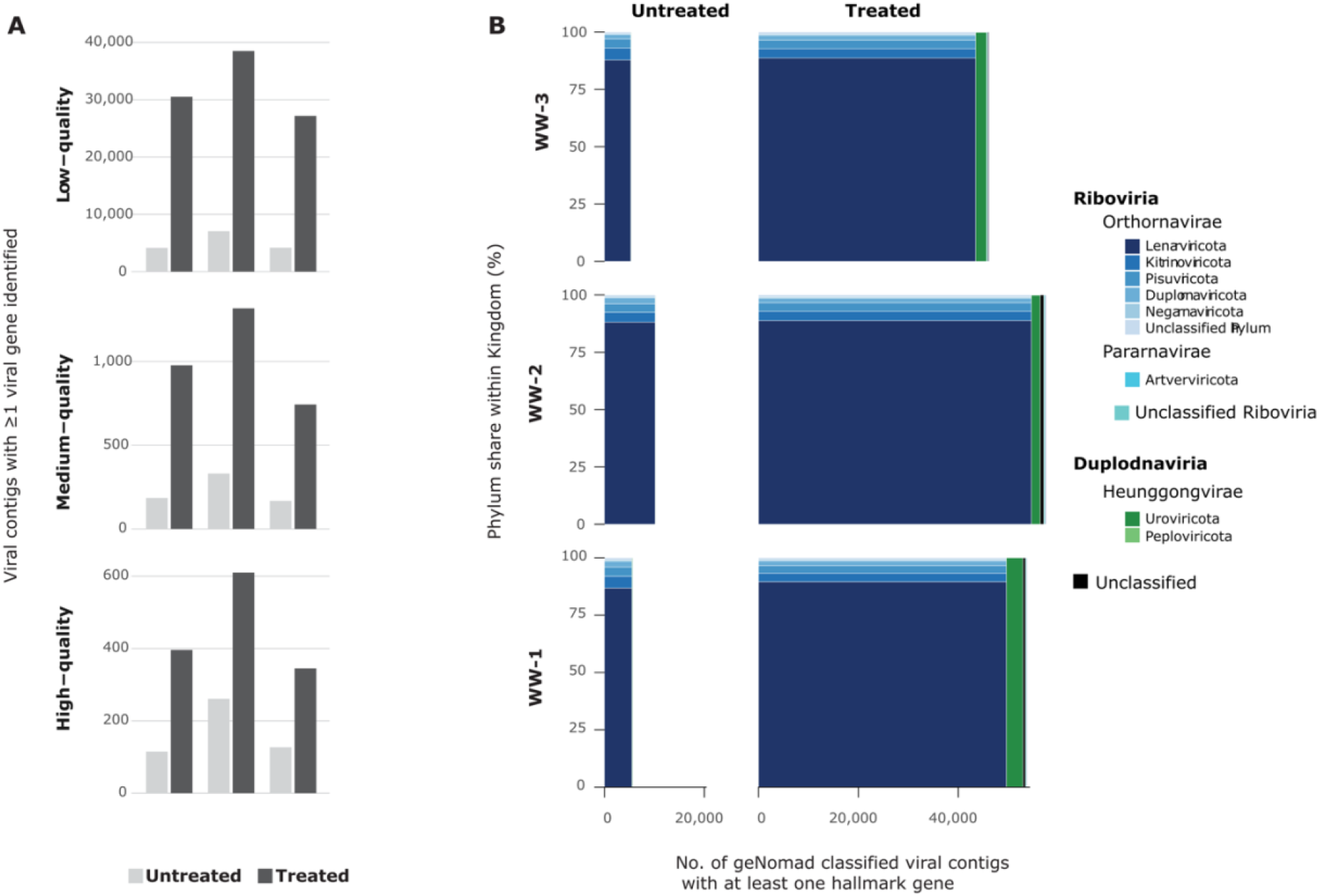
Targeted depletion increases assembled viral contigs in wastewater metatranscriptomes. (A) Number of assembled viral contigs per library with ≥1 viral gene identified by CheckV. (B) Number of geNomad classified viral contigs per library with ≥1 viral hallmark gene. The width of each column represents the total number of viral contigs within each kingdom (Riboviria or Duplodnaviria) and the y-axis represents the percentage of each phylum within each kingdom.

Quality assessment of viral contigs using CheckV showed an increase in low-, medium-, and high-quality contigs containing at least one viral gene (Figure 2A and Supplementary Table 4). Across all samples, the majority of viral contigs were categorized as low-quality, which is expected for metatranscriptomic data, where assemblies can represent expressed genomic fragments rather than complete viral genomes. However, depletion-treated samples showed enhanced viral recovery across all quality tiers, including medium- and high-quality viral contigs, indicating improved viral representation in terms of both quantity and quality.

### Improved detection of human and animal pathogens after depletion treatment

To determine how the depletion treatment impacted the detection and recovery of potential public health pathogens, non-rRNA reads were aligned to a curated human/animal/plant viral pathogen database using EsViritu. Proportions of total reads aligned to the viral database in treated samples increased on average 42-fold compared to untreated samples, accounting for between 168,936 and 326,643 additional viral pathogen reads. This represented between 0.02% and 0.04% of total sequenced reads (Figure 3A, Supplementary Table 5). Furthermore, 161 EsViritu-classified viral species were identified on average per treated sample compared to 71 in the untreated samples, more than doubling the number of detected viral species (Figure 3B). Importantly, several human pathogens of public health concern, including influenza A, B, and C, hepatitis A, rhinovirus B, respiratory syncytial virus (RSV), human metapneumovirus, human parainfluenza virus 4, and astrovirus VA3, were detected in treated samples but were undetected in the untreated samples (Figure 3C, Supplementary Table 5). Among the viral species detected in both treated and untreated samples, species in the treated metatranscriptomes exhibited a 33-fold increase in assigned read counts on average (Figure 3D), providing increased genomic depth for improved variant calling and phylogenetic tracking. Overall, RNA depletion substantially increased the diversity, abundance, and detection of pathogenic viruses from wastewater, enabling improved characterization of viral strains that were not detectable in untreated samples.

**Figure 3.**
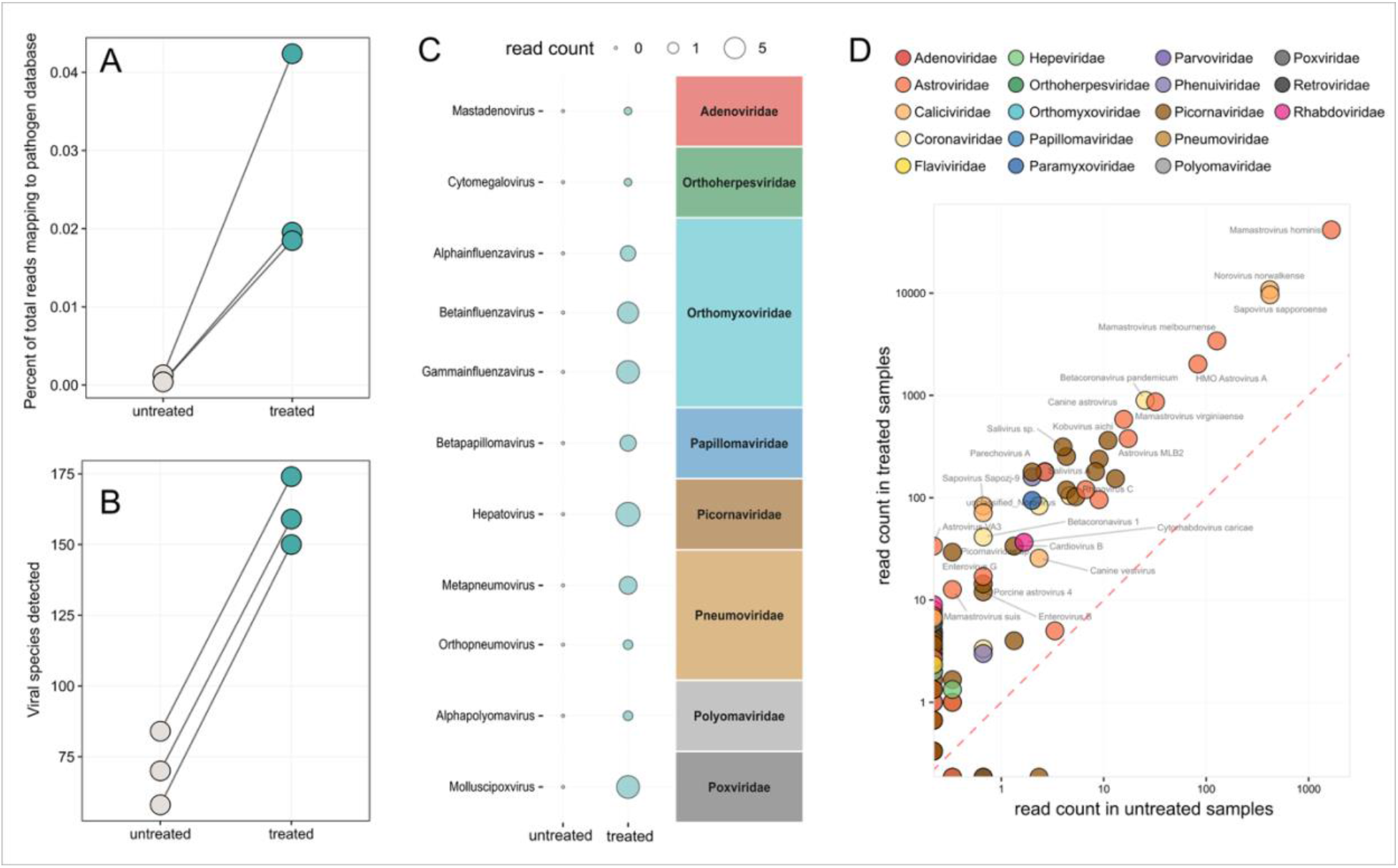
Increased detection of pathogens in treated samples. A) Proportion of reads mapping to the Viral Pathogen Database in treated and untreated samples. Lines connect paired samples. B) Number of viral species detected in treated and untreated samples. Lines connect paired samples. C) Average read count of genera from human-relevant viral families detected in treated samples but not in untreated samples. D) Average read counts of detected viral species from human-relevant viral families in treated and untreated samples. Red dashed line shows a 1:1 ratio. Due to the log-transform, viral species that were absent in the untreated and treated samples are located along the y- and x-axes respectively.

## Discussion

Tobamoviruses have repeatedly been reported among the most abundant RNA viruses in wastewater metatranscriptomes worldwide^26,27,37^. These plant viruses infect food crops, particularly peppers and tomatoes, and are consumed in fresh produce and processed foods such as pepper-based sauces and seasonings. The stable viral particles can survive passage through the gastrointestinal tract and are shed in feces, providing a relatively constant dietary source of tobamovirus RNA in wastewater ^38–40^. While potentially useful in other contexts (e.g., as fecal normalization markers)^41–43^, in the context of sequencing, this signal is less helpful, occupying up to 25% of viral reads in our untreated samples. The depletion strategy used here lowered this undesired signal to <2% using a customized probe panel designed for this study. Accordingly, reads assigned to other RNA viruses (Riboviria) increased substantially, with non-tobamovirus signals representing up to 69% of viral reads and an increase in the number of respective viral contigs in treated libraries. Together, these findings indicate that targeted depletion probes can redirect sequencing effort away from abundantly represented taxa and improve the capacity to study RNA viral diversity.

Depletion of rRNA was also successful as the vast majority of reads (>90%) in untreated samples were rRNAs while treated samples contained <5% rRNA. Although rRNA removal is common in RNA-Seq experiments, it is important to note its utility in viral wastewater metatranscriptomics. In the wastewater matrix, bacterial and eukaryotic cells can lyse during sample transit and processing, releasing abundant ribosomal RNAs into the supernatant. These rRNA molecules can be concentrated with viral particles during ultrafiltration steps, and will be sequenced alongside target genomes from RNA viruses. The large increase in non-rRNA reads after the depletion treatment supports including rRNA removal as a standard step in wastewater RNA sequencing workflows, especially when procedures to remove unencapsidated nucleic acids prior to extraction have not been performed.

The depletion probes caused the average proportion of viral pathogen reads to increase by 42 fold, with upwards of 165,000 additional viral pathogen reads available in treated samples. The additional viral reads uncovered following depletion included signals from clinically relevant pathogens like influenza, RSV, and hepatitis A, which were missed in the untreated samples. The latter example is especially impressive, given how rare hepatitis A virus is in Calgary wastewater ^9^ compared to influenza and RSV which are detected with higher frequency ^44^. Although pathogenic viral sequences represented a relatively small proportion (0.02–0.04%) of the total genomic content in the treated wastewater metatranscriptomes, these proportions approached those typically observed following pathogen enrichment using capture probes targeting thousands of viruses ^8,19,20^.

The sample processing strategy used here was designed to recover RNA from concentrated wastewater supernatants, and likely also captured extracellular nucleic acids and small non-viral particles in addition to intact virions. More stringent particle purification approaches, such as nuclease treatment of concentrated samples prior to extraction^27^ or alternative concentration methods such as PEG precipitation^28^, may further enrich viral nucleic acids and reduce background signals derived from lysed cells. Combined with the depletion strategy presented here, such steps could improve viral read output further. On the other hand, nuclease pre-treatment might also degrade nucleic acids from viral particles of interest that are not intact. In wastewater, non-intact viral particles are not uncommon following partial degradation within the human host and during transit through the sewer system. Genomic sequences from DNA viruses could also be detected in the three wastewater datasets analyzed here, despite extensive DNase treatment after nucleic acid extraction, further demonstrating that non-target nucleic acids are not fully eliminated during sample processing. This was a persistent feature after several different attempts on various samples and using different enzymes.

For individual tobamovirus targets, depletion was successful for seven of the eight candidates included in the probe panel. The only exception was Tobacco Mild Green Mosaic Virus, for which read abundance did not decrease. However, based on the untreated metatranscriptome data, this virus was present at relatively low abundance in the untreated samples compared with the other seven targets (Figure 1C), therefore the lack of successful depletion using this probe did not substantially affect the overall success of the tobamovirus removal strategy. The lack of depletion may reflect a sequence mismatch between particular environmental strains of Tobacco Mild Green Mosaic Virus present in Calgary wastewater samples and the reference sequence used for probe design. Because these probes are highly sequence specific and have limited tolerance for mismatches, even modest sequence divergence could reduce hybridization efficiency. In addition, probe selection prioritized the most abundant viruses, while the number of probes targeting lower-abundance viruses and sequence variants was minimized to maintain a reasonable overall panel cost.

Benefits of depletion extended beyond read-based analyses and were also evident in assembly-based recovery of viral contigs. Depleted libraries yielded substantially more viral contigs as classified by both CheckV and geNomad, including increases in low-, medium-, and high-quality viral sequences. This is a meaningful outcome since improved recovery of assembled viral genomes enhances the potential for genome-resolved analyses and discovery of novel viruses. In particular, geNomad identified a marked increase in viral contigs containing hallmark genes for RNA viruses that this study specifically undertook to resolve.

Most of the additional viral contigs recovered after depletion belong to the phylum Lenarviricota, particularly to the class Leviviricetes, although many could not be classified beyond class level. Leviviricetes comprises single-stranded RNA bacteriophages that are widely distributed in animal and human feces and thus also municipal wastewater ^45–47^. The large number of contigs that remained unclassified at finer taxonomic levels suggests that wastewater contains substantial, previously undescribed RNA viral diversity. RNA bacteriophages remain comparatively understudied relative to their DNA counterparts, despite their widespread occurrence in wastewater and other microbial communities, like the human gut ^48,49^. Consequently, their function in the environment, including their potential influence on the human microbiome and horizontal gene transfer, remain poorly understood. Many Leviviricetes contigs were around 4,000 bp in length and may represent near-complete or complete genomes of previously uncharacterized viruses. Together, these findings demonstrate that targeted depletion of dominant wastewater RNA sequences can improve access to otherwise poorly represented components of the wastewater virome, including understudied RNA bacteriophages.

In future studies, subtractive depletion could be combined with ultra-deep untargeted sequencing to improve pathogen detection efficiency. Previously, untargeted sequencing at approximately one billion reads per sample enabled detection of expected and unexpected human pathogens, however they found that plant viruses such as tobamoviruses and rRNA sequences were highly abundant ^18,23^. These findings reinforce that sequencing depth alone does not fully address the low target-to-background ratio characteristic of wastewater. Our study shows that combining untargeted sequencing with depletion of consistently abundant, well-characterized background RNAs redirects a greater fraction of sequencing capacity toward lower-abundance sequences and improves the probability that rare viral genomes receive sufficient coverage for detection and assembly. The same principle could be extended to target other abundant baseline viruses for depletion. Such panels would require careful design so that removal of background taxa does not compromise surveillance objectives, particularly where abundant plant viruses or bacteriophages are themselves of interest.

In conclusion, combined depletion of abundant rRNA and tobamovirus sequences substantially increased the sequencing capacity available for other wastewater viruses, improving the detection of human pathogens and the recovery of contigs from poorly characterized RNA viruses. Importantly, this approach enhances the efficiency of untargeted wastewater metatranscriptomics while preserving its capacity to detect a broad range of viral taxa beyond those included in predefined enrichment panels. By reducing competition from highly abundant background sequences, targeted depletion offers a practical strategy for making wastewater sequencing more informative and expanding its potential for pathogen surveillance and viral discovery.

## Acknowledgements

We thank Francisco Daniel Dávila Alemán for helpful discussions and brainstorming related to data visualization and figure development.

